# Taking sharper pictures of malaria with CAMERAs: Combined Antibodies to Measure Exposure Recency Assays

**DOI:** 10.1101/299446

**Authors:** Bryan Greenhouse, David L. Smith, Isabel Rodríguez-Barraquer, Ivo Mueller, Chris J. Drakeley

## Abstract

Antibodies directed against malaria parasites are easy and inexpensive to measure but remain an underutilized surveillance tool due to a lack of consensus on what to measure and how to interpret results. High throughput screening of antibodies from well-characterized cohorts offers a means to substantially improve existing assays by rationally choosing the most informative sets of responses and analytical methods. Recent data suggest that high-resolution data on malaria exposure can be obtained from a small number of samples by measuring a handful of properly chosen antibody responses. In this review, we will discuss how standardized multi-antibody assays can be developed and efficiently integrated into existing surveillance activities, with great potential to greatly augment the breadth and quality of information available to direct and monitor malaria control and elimination efforts.

## Effective malaria strategies require good data

Elimination of malaria or effective and sustainable control require deployment of interventions updated using surveillance data. Contemporary, accurate information on malaria transmission is required to determine which interventions should be deployed where to reduce transmission effectively given resource constraints. The ability to detect changes in transmission is an integral part of evaluating programe activities by providing evidence for the effectiveness of interventions and identifying when and where changes are needed. Given the central role of surveillance data in guiding strategy, any tools that could provide data more accurately or cost-effectively would be of great value. The next generation of antibody assays for measuring malaria transmission holds promise for doing all three.

## The current focus of malaria surveillance

One would be hard-pressed to find a malaria program officer, epidemiologist, or funder of interventions who would not prefer a clearer picture of malaria in their target population. Some types of surveillance data, such as detailed entomological measurements or cohort studies, are not feasible for routine surveillance of transmission since they are too resource intensive to be performed extensively. These infrequent high-resolution snapshots are extremely valuable for answering research questions but do not provide a picture of malaria transmission complete enough for programmatic use. The rest of the picture is filled in primarily from two other sources of data – routine clinical surveillance and cross-sectional surveys. Clinical surveillance data are widely and continuously collected as health systems provide care to those with malaria. The utility of these data depend heavily on the quality of diagnosis, completeness of reporting, and ability to account for important factors regarding catchment populations, care seeking, and clinical immunity ^1^. Thus, while the quality of and access to these data are improving, limitations remain, particularly in settings where a small minority of infections get reported through standard surveillance ^2^.

The most widely collected, standardized data currently available are based on surveys that collect blood from a cross-sectional sample of individuals in a population. The primary metric collected in most surveys to date has been the prevalence of parasites or parasite rate (PR), based on light microscopy, rapid diagnostic test (RDT), or nucleic acid detection (e.g. PCR). Data from such surveys form the basis of global maps of malaria ^3^. Prevalence data, while useful, are limited in the ability to detect changes in malaria transmission where transmission is so high that PR remains high even if exposure decreases considerably, or where it is so low that infeasibly large sample sizes are required to accurately measure changes over time or at fine enough spatial scales to be programmatically useful ^4^.

Prevalence data are an important mainstay of surveillance, but each sample provides a single piece of information – whether a person has detectable blood stage infection or not. This is a particular challenge in seasonal transmission environments or with *P. vivax*, where people can harbor dormant liver stage infection without having a concurrent bloodstage infection. It is possible to learn more about malaria exposure from the same blood sample by additionally measuring antibody responses to parasite antigens. This holds the promise that a single sample could add substantially more information to population level measures of transmission, since each antibody response measured provides information about past exposure in addition to current infection status. Obtaining antibody data is quite practical: ELISA and some multiplexed assays are quite inexpensive and can be performed on material extracted from dried blood spots, which are simple to collect and transport; point-of-contact tests such as lateral flow and microfluidic assays are also options. The result promises to be a more resolved digital picture of malaria transmission.

## Using antibodies to sharpen surveillance data

There is growing interest in using serosurveys to gain understanding of disease transmission and inform control interventions across a broad set of pathogens ^5^. Antibodies have been used to estimate exposure to pathogens, including malaria parasites for over 70 years, but the approach has become more accessible and standardized due to availability of purified recombinant antigens and development of appropriate analytical methods. A commonly used strategy to estimate transmission intensity from serological surveys has been to analyze sero-positivity by age to compute a force of infection or seroconversion rate (SCR), which takes advantage of the fact that individuals in more highly endemic areas are more likely to have been exposed to malaria parasites and thus to have detectable antibody responses. These methods were initially developed for permanently immunizing infections such as measles and yellow fever ^6,7^, but adapted for malaria in the last decade ^8,9^. By translating age-stratified antibody prevalence data into a metric which reflects overall transmission in a community, the SCR extracts meaningful population information from individuals’ binary, typically long-lived responses to immunogenic Plasmodium proteins (Figure 1A). This strategy has been validated across a wide range of epidemiologic settings, and clearly demonstrates the value of collecting antibody data in surveys (Figure 1B).

**Figure 1:**
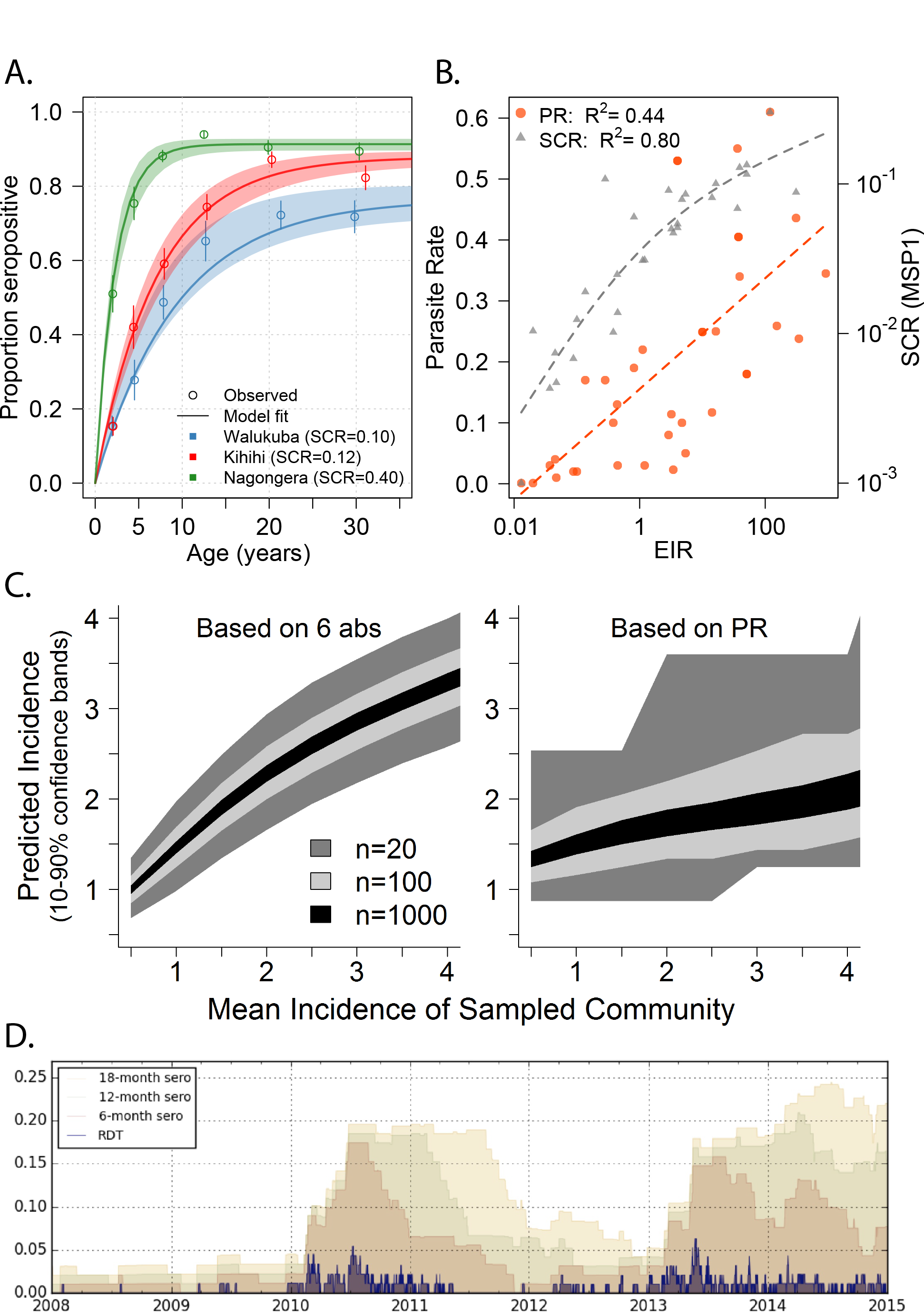
Established and next-generation methods for evaluating malaria transmission via antibodies provide higher resolution than parasite prevalence. A. The seroconversion rate (SCR) for a population can be calculated from age-stratified prevalence of antibody responses, often with a long half-life. Data shown here are responses to apical membrane antigen 1 (AMA-1) from 3 cross-sectionals surveys in Uganda ^25^. B. Paired SCR and parasite rate (PR) data from multiple sites ^10,25–34^ demonstrate that SCR (using merozoite surface protein 1, MSP-1) has a tighter association with transmission, as measured by the annual entomologic inoculation rate (EIR). C. Using 6 antibodies identified as informative about recent exposure, predictions of Pf exposure in a community can be obtained from relatively small surveys, in contrast to PR data obtained from the same surveys ^19^. D) A simulation of a small village (n=100) with seasonal, low transmission illustrates how ongoing transmission can be detected consistently from an antibody test measuring recent exposure, but less reliably from RDT ^35^.

The SCR provides a useful metric for stable, community-level transmission, but with recent successes in decreasing malaria transmission it is frequently of interest to measure changes in exposure over time. Signals of abrupt changes in exposure have been detected in the age-stratified antibody prevalence profile by estimating a change in SCR, reflecting a change transmission intensity at some point in time or with a certain age ^10^. However, it is difficult to quantify very recent changes in exposure from binary antibody responses with a long half-life ^11^. Similarly, it is often difficult to quantify changes in transmission from samples collected at a single time-point as they may be confounded by age dependence in risk. Fortunately, the human antibody response to complex pathogens contains rich information which can be leveraged if the right responses are identified, measured, and analyzed appropriately. Advances are being made in three related areas: 1) using information contained in the titers of antibodies, instead of reducing them to binary responses; 2) measuring antibodies to multiple antigens with differing kinetics, rather than limiting measurement to one or a few antigens with long half-lives; and 3) developing analytical methods which take advantage of these rich data to provide precise, quantitative estimates of exposure history in populations from more intensive analysis of fewer samples.

Methods have been recently developed to extend the analysis of age stratified antibody prevalence by incorporating antibody titers ^12–14^. Some of these models are similar in concept to those used to calculate SCR, but allow for different rates of boost and decay of relative antibody concentrations - data which are often already available from standardized ELISA or multiplex assay readouts. By using more information, these methods should produce more precise estimates of malaria transmission than those derived from binary responses. In parallel, investigators have begun to evaluate what information can be obtained by measuring responses to different antigens ^15–18^. Not surprisingly, responses to different antigens appear to provide information of greater or lesser value for answering specific epidemiologic questions in different populations depending on their immunogenicity and other properties. For example, by evaluating SCR to hundreds of proteins using a protein microarray, Baum et al showed that a distinct subset of these targets were more efficient in distinguishing transmission in two areas of Highland Kenya ^15^. Similarly, Ondigo et al demonstrated that seroconversion and reversion rates to different antigens varied considerably, and illustrated how evaluating SCR to a number of antigens could provide more information on the temporal structure of past exposure than looking at responses to a single antigen ^16^. To turn these promising findings into reliable, informative antibody assays, we propose a strategy to design Combined Antibodies to Measure Exposure Recency Assays (CAMERAs) with the aim of answering actionable questions across a range of epidemiologic settings ^19^.

## Designing high resolution CAMERAs for actionable malaria surveillance

Obtaining precision surveillance data from antibodies requires a detailed understanding of how antibody responses change over time in response to infection, and how these kinetics are modulated by factors such as age, genetic diversity in parasite and host, and prior Plasmodium exposure (including different species). Initiating prospective studies to define antibody kinetics throughout a representative set of epidemiologic settings would be an enormous endeavor.

Fortunately, a number of well-characterized cohorts have been performed in malaria endemic areas, and many have archived appropriate biologic material in the form of serum, plasma, or dried blood spots. With these existing studies providing at least a subset of the samples needed, the next step is to identify antibody responses that are most informative about transmission. While it has been shown that different antigens tend to elicit different qualities of response ^20,21^, it is still difficult to predict these a priori. In addition, it is not just the average of any particular metric of response to an antigen (e.g. degree of boosting with exposure, half-life, etc.) that matters but also biological variation in these qualities across individuals. Therefore, a broad screen of responses provides the highest probability of identifying the optimal set of informative biomarkers.

High throughput screens using protein microarrays or similar technologies offer one such approach ^18,22^. This technique allows rapid screening of a large number of responses and requires minimal amounts of both sample and antigen. Rapid production of “crude” antigens without much optimization allows a large number of antibody responses to be screened up front with minimal start up time. One potential downside of this approach is that a subset of antigens may not be properly expressed or folded, but this potential limitation is less of an issue for efforts trying to identify markers of exposure as opposed to characterizing correlates of immunity. Furthermore this downside is potentially overcome by a numbers game – responses to certain sets of antigens are likely to provide similar information, and by screening many analytes simultaneously it is likely that at least one representative member of each set will be successfully evaluated. Other high throughput approaches, including peptide arrays ^23^ and phage display libraries ^24^, may allow rapid evaluation of individual linear epitopes, including the ability to capture naturally occurring genetic variation in the target. The effort in producing standardized reagents at scale – ultimately required for final assays – can then be focused on a down selected set of promising antigens followed by iterations of validation to design CAMERAs (Figure 2).

**Figure 2:**
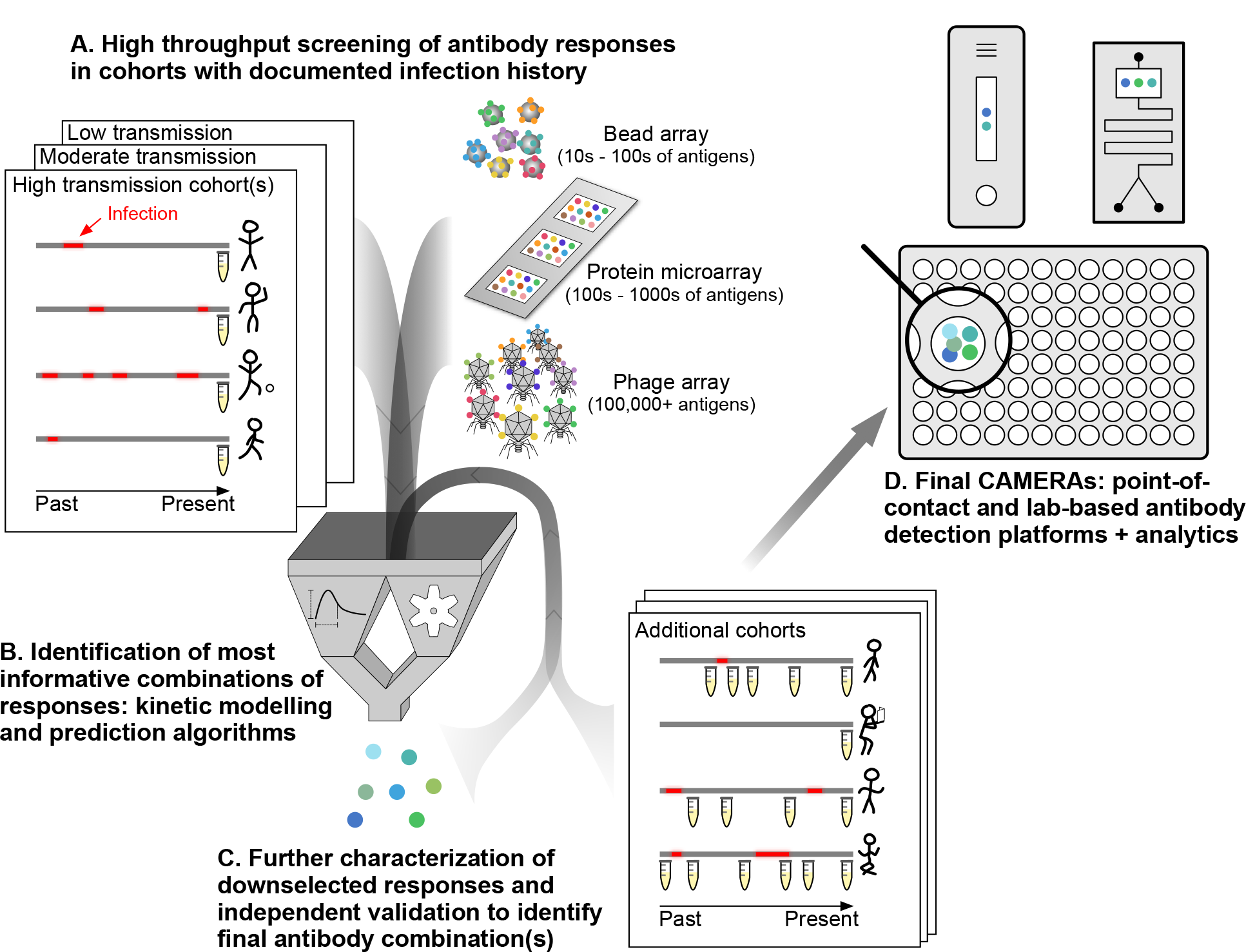
Approach to designing Combined Antibodies to Measure Exposure Recency Assays (CAMERAs) A) Samples from detailed cohorts, where accurate data on individuals’ prior malaria infections are available, are critical for providing a gold standard to identify informative antibody responses. Cohorts should represent the range of ages and epidemiologic settings where CAMERAs will ultimately be used. Various platforms are available for high throughput screening of antibody responses, with tradeoffs based on cost, number of analytes that can be screened, precision, and dynamic range. B) Downselection of the most informative combinations of responses (i.e. considered jointly) is accomplished via parametric modeling of antibody kinetics ^36^ and/or any number of machine learning prediction algorithms. Both of these analytical approaches have advantages, and combining both may be optimal. C) Top “hits” identified in comprehensive screens require validation in distinct individuals and cohorts. Given the smaller number of responses evaluated, it may be feasible to evaluate much larger numbers of samples including longitudinal sampling from individuals over time. D) Final CAMERAs can be designed as point-of-contact (e.g. based on lateral flow or microfluidics) or laboratory based assays, depending on the use case. The analytics for deriving epidemiologically relevant metrics from antibody responses will be integral to the assay.

We have recently demonstrated the feasibility and potential of this approach, screening plasma samples from two cohorts of children in Uganda using a protein microarray containing 856 *P. falciparum* antigens included based on their potential to provide information about prior exposure to this parasite ^19^. In this study, data from a small subset of antibodies (e.g. 3-6) chosen based on their combined information content measured at a single timepoint provided accurate estimates of whether or not an individual was infected in the last 30, 90, or 365 days and on their incidence of malaria in the prior year. These quantitative, individual-level data rivaled the rich information obtained from expensive cohort studies. For example, estimates of incidence obtained from antibodies accurately reproduced the spatial heterogeneity in transmission within a community detected by one year of entomological and clinical cohort data. When aggregated, antibody data from individuals dramatically outperformed parasite prevalence in precision and dynamic range, providing estimates of the incidence of malaria in a community from sample sizes as small as 20 (Figure 1C).

The specific antibody responses selected to be measured will depend on which are most informative for a given scenario. The number of potential antigens encoded by Plasmodium genomes, and number of sequence variants for those under balancing selection, is large, providing an embarrassment of riches regarding what to measure. Antibodies directed at antigens expressed exclusively during the pre-erythrocytic stage of infection may make good markers of newly acquired infection; those continually boosted by low-density asexual stage infection may be better indicators of recent chronic infection; those specific for a given Plasmodia will provide species-specific data. Evaluating responses to a panel of antigenic variants, or even specific epitopes within these variants, looking at different classes or IgG subclasses of antibodies, and evaluating avidity may all in theory provide additional information given the different kinetics of these responses. Antibodies directed against mosquito salivary antigens may provide additional information regarding exposure to important vectors. In the end, consistent empiric data validated across multiple settings with appropriate gold standards will be the best arbiter of what should be included in a CAMERA. A number of platforms are currently available for measuring antibodies (Figure 2), and options will continue to grow. Important factors informing appropriate platforms include the number and type of antibody responses and whether binary, semi-quantitative, or quantitative responses are required; cost, portability, speed, and level of training and equipment needed to perform the assay.

With the availability of samples and data from cohorts, methods for high throughput screening of antibody responses, and a robust data analysis approach considering combinations of responses with appropriate validation, it should be possible to craft antibody assays that will provide accurate answers to nearly any malaria surveillance question with respect to the rate and timing of human Plasmodium infection. With that in mind, the most pressing questions will largely depend on the epidemiologic setting and consumer of the data (Table 1). In areas of endemic transmission, it will be important to measure variations in transmission over space and time, especially in response to interventions. For control programs deciding strategy in areas of very low transmission attempting to eliminate malaria, additional questions become important. In these areas, data on recent infections are of particular added value, given the large sample sizes otherwise needed to confirm or exclude ongoing transmission (Figure 1D). Finally, in the research setting, biomarkers of individuals’ prior malaria exposure may be important outcomes in epidemiologic studies, or as ways to adjust analyses for heterogeneous exposure, e.g. when investigating mechanisms of immunologic protection.

**Table 1.** Actionable malaria surveillance data obtainable with CAMERAs

**Figure 3. Outstanding questions for developing and using CAMERAs.**
| Setting | Relevant questions | Information derived from antibody assays | Added value to traditional metrics |
| --- | --- | --- | --- |
| Programmatic, endemic | What is the current level of transmission and how does it vary over space and time? | Accurate estimates of transmission intensity calibrated to relevant metrics (e.g. force of infection) now and over time for communities. In particular, how intensity changes in response to interventions, human and mosquito behavior, and other factors. | Dynamic range allows estimates over a broader range of transmission intensity than parasite prevalence. |
|  | Where are interventions most needed, and which are optimal? | Estimates of recent *P. vivax* infection, and thus likely hypnozoite carriage. | Increased precision allows for smaller sample sizes and/or spatial mapping at a more granular level. |
|  | How well are current interventions working? |  | Ability to measure prior as well as current transmission, for areas where prior estimates are not available. |
|  | Are individuals infected with dormant stages of parasites (*P. vivax*)? |  | Ability to determine whether individuals are latently infected with *P. vivax*. |
| Programmatic, peri-elimination | (above, plus) | Where recently or currently infected individuals live. | (above plus) |
|  | Where are residual foci of transmission, if any? | Identification of parasite species causing recent infections. | More information from each individual allows for smaller sample sizes and/or more granular spatial data. |
|  | Which Plasmodium species are causing infections? | Demographics of recently or currently infected individuals. | Increased sensitivity for detecting infections when they are rare, including by species such as *P. knowlesi*, by detecting over a larger range of time. |
|  | What are the demographic groups at highest risk of infection or transmission to others? | How far in the past infections took place. |  |
|  | Has transmission been interrupted? | Historical spatial distribution of malaria exposure. | Ability to reconstruct historical exposure from contemporary measurements. |
|  | What is the receptivity of the area? | Probability of individuals experiencing symptomatic or severe | Ability to measure waning immunity. |
|  | If the population |  |  |

|  | susceptible to epidemic transmission? | disease upon infection. |  |
| --- | --- | --- | --- |
| Research | What are the epidemiologic risk factors for infection with malaria parasites? | Estimates of individuals' prior exposure.<br>Determining how much variation in naturally acquired immunity can be attributed to differences in prior exposure. | Ability to evaluate diversity of parasites to which individuals have been previously exposed, e.g. by measuring breadth of responses to polymorphic antigens.<br>Ability to estimate individuals' cumulative and recent exposure prior to observation during the research study. |
|  | What are the biomarkers and mechanisms of immunity to malaria? |  |  |

## People are different, and that’s ok

The extent to which particular surveillance questions can be answered by antibody data, and the corresponding sample sizes required to obtain answers with a given accuracy, will be largely determined by the sources of variation in antibody responses and the degree to which this variability can be systematically accounted for. Whilst laboratory methods can be optimized to minimize technical variation in the measurement of titer or presence vs. absence of an antibody response, biological variation will remain. Some of this biological variation may be attributable to identifiable factors such as age and history of prior infection, but some will be unmeasured biological variability that is difficult to account for, e.g. due to host genetics or nutritional status. Despite this variation, precise population level estimates can be obtained from antibody responses by 1) measuring responses to multiple antigens; 2) sampling multiple people in a given area; 3) incorporating knowledge of age-exposure-antibody relationships into data analysis; and 4) tailoring antibody assays to specific age groups and/or transmission intensities. Regarding the latter, while having a universal antibody assay for use in all settings may seem the most straightforward, in reality it is likely that certain sets of responses will have more utility in some settings versus others. For example, antibody responses which provide information about a decline in EIR from 100 to 10 infectious bites per person year may be different than those which can confirm absence of recent exposure in an elimination setting. Depending on multiplex capacity, the same standardized laboratory platform may essentially contain multiple “assays”, of which only a subset are actually used to produce outputs for a given setting. Platforms requiring a more parsimonious set of analytes, e.g. standard lateral flow assays, may be best targeted to a specific epidemiologic setting.

The apparent ambiguity of how to interpret a particular antibody response in a particular individual has been a psychological barrier in the widespread dissemination and acceptance of antibody data for surveillance. However, more commonly used metrics, such as parasite prevalence, are subject to similar caveats. Detecting a parasite in an individual’s blood, while seemingly concrete, is still an indirect measure of transmission mediated by the duration and density of infection, which are functions of age, prior exposure, and the limit of detection of the assay used (e.g. microscopy vs. PCR). For this reason, parasite prevalence requires age standardization a reasonably sized sample of the population to obtain a single point estimate ^3^. Estimates of transmission derived from antibodies will also be averaged across a population sample. Thus, while there will be some uncertainty as to, e.g., if a particular individual has been infected 2 vs. 3 times in the past year, precise estimates of transmission can still be derived for the population. As discussed earlier, the increased granularity of information provided by antibody data for each individual will likely result in more accurate estimates from smaller sample sizes despite inherent biological variation. This advantage holds across all transmission intensities, but may be particularly salient in areas of very low transmission.

## Practical considerations for incorporating CAMERAs into routine surveillance

Antibody measurements have great potential for augmenting information obtained in malaria surveillance activities. However, specific questions remain on what should be measured, in whom, and how to best make sense of the data (Figure 3). This will inevitably be an iterative process, with analytes, platforms, and analytical methods building on prior efforts. If methods are designed to map to relevant metrics of interest, e.g. force of infection in the last year or presence or absence of infection within the past year, then methods with improved test characteristics can be implemented as they become available with comparable outputs.

**Figure 3:**
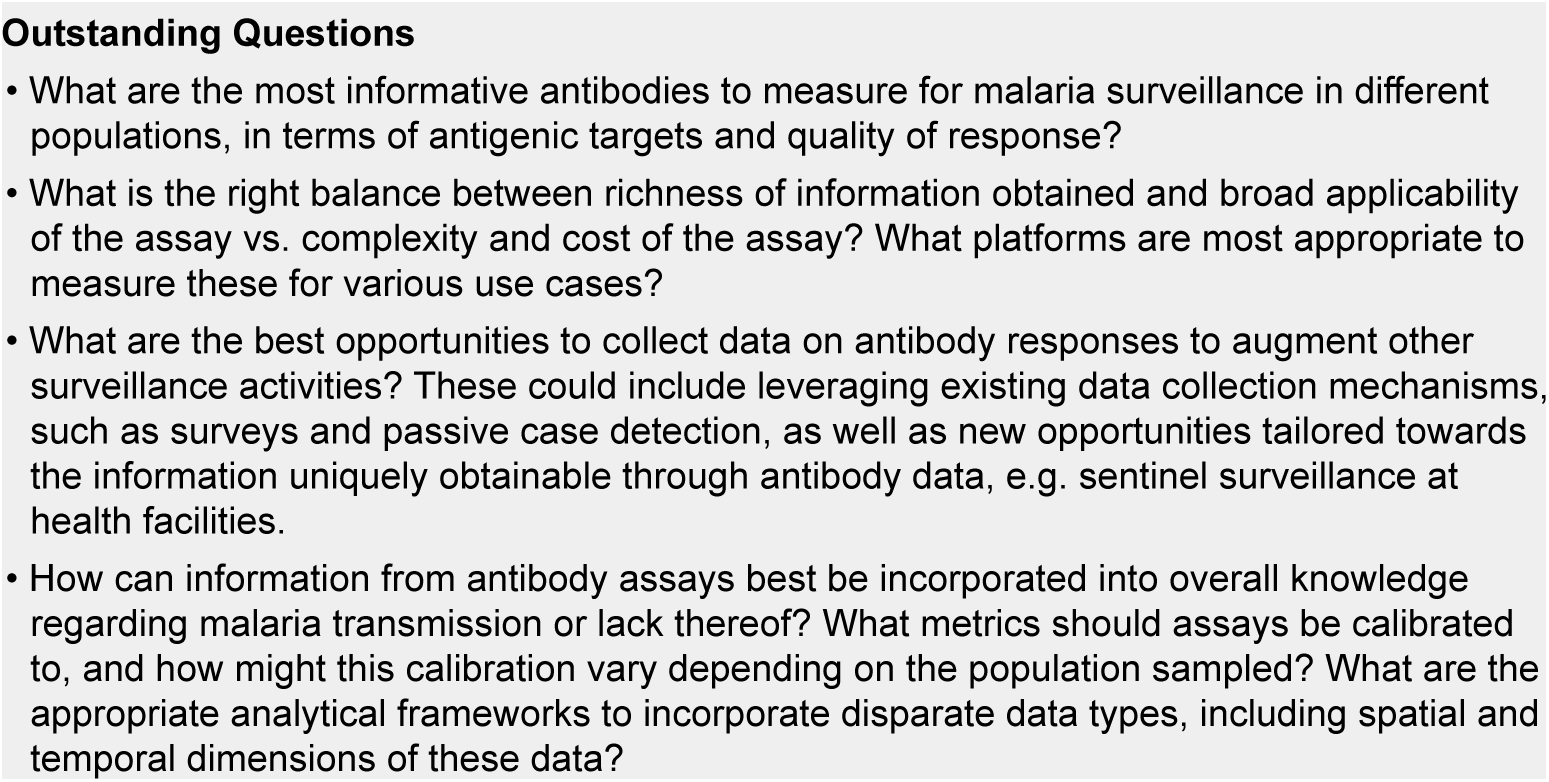
Outstanding questions for developing and using CAMERAs

Once CAMERAs are designed and test characteristics are known, e.g. amongst which age groups they are informative in a given setting, appropriate surveillance activities for sample collection and antibody measurement can be determined. For cross-sectional surveys, e.g. malaria indicator surveys, school surveys, and other surveillance activities including those performed in intervention trials, CAMERAs offer a clear opportunity for obtaining additional information. Such measurements will add little incremental cost, and in fact may offer substantial cost savings as smaller sample sizes may be required to obtain similar information. Incorporating measurement of antibodies in many cases requires no additional field work, since many such surveys are already collecting dried blood spot samples. For example, DHS surveys conducted in over 50 countries already collect serum samples to measure specific biomarkers ^5^. Efficient collection of samples from other convenient venues, e.g. from those presenting to health facilities for routine care, may provide continuous, low-cost data to augment metrics such as test positivity rates and malaria incidence. The ease of collecting more comprehensive data at low cost via such approaches will need to be balanced with issues surrounding potentially biased sampling, and is an area ripe for further investigation.

Finally, antibody data generated need to be consistently translated into easily interpretable metrics of transmission. Development and evaluation of analytical methods will be an integral part of assay design, critical for making decisions regarding the number and type of antibody responses which will be measured. Simple point of care assays may require straightforward interpretation, e.g. a band visible on a lateral flow assay might indicate infection within the past year. However, given the potential increase in the breadth and accuracy of information obtained from combining data from multiple antibody responses, other assays may utilize more sophisticated algorithms. Such algorithms can be easily implemented in software to provide straightforward interpretation regardless of the complexity of the underlying algorithm.

The path for developing high resolution CAMERAs is clear, and a number of research teams are working toward answering the salient questions outlined above. As CAMERAs are designed, validated, and improved, their key role in malaria surveillance will come into clear focus.

## Acknowledgements

We thank Edward Wenger (Institude for Disease Modeling) for producing the simulation and associated image used in Figure 1D. This work was funded in part by the NIH/NIAID International Centers of Excellence in Malaria Research (ICEMR) program, grants U19 AI089674 and U19 AI129392, NIH/NIAID R01 AI119019, and National Health & Medical Research Fund (GNT1092789). BG is a Chan Zuckerberg Biohub investigator.

